## Supplemental Figures for "A novel interface for cortical columnar neuromodulation with multi-point infrared neural stimulation"

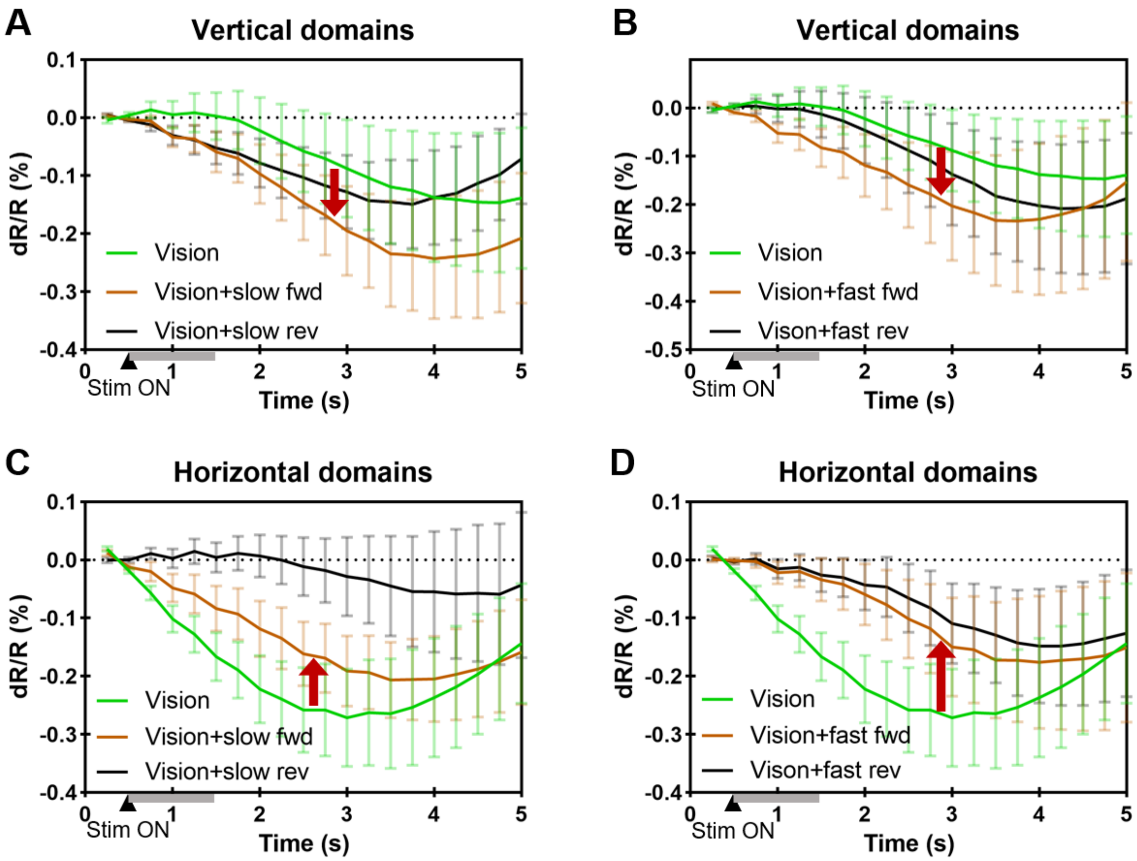


Supplementary Fig.1. Directionality. OI Response for ROIs in right panel of Fig. 5A on right hemisphere during different directions of 'vertical INS' on left hemisphere. (A & B) 2 directions with slow or fast speed, both enhance the vertical visual response (INS and visual stimulation are matched). (C & D) 2 directions with slow or fast speed, both reduce the horizontal visual response (INS and visual stimulation are non-matched). FWD: posterior to anterior, away from HM. REV: anterior to posterior, towards HM. All symbols are same as Figure 5.

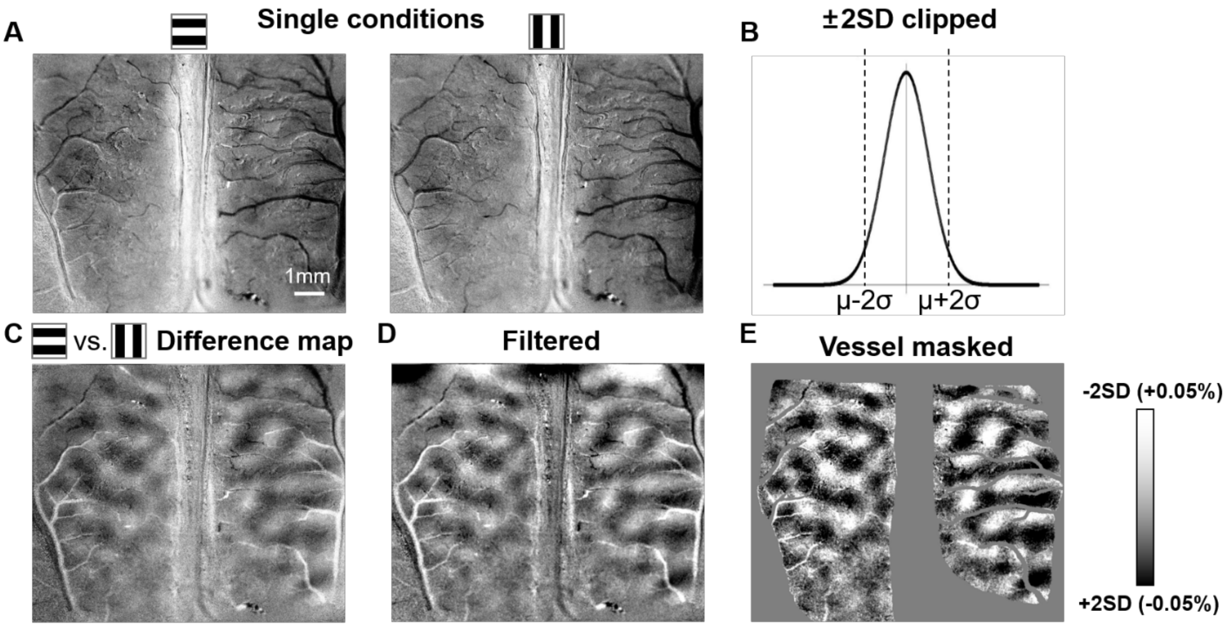


Supplementary Fig.2. Steps in processing of OI data. (A) Single condition maps or horizontal (left) and vertical (right) gratings. (B) In the grayscale distribution of an image, we 'clip' the distribution to remove large artifactual (e.g. dR/R > 2%) reflectances which are often due to large vascular pulsations or locations of specularity. A clip of median ± 2SD is typical. (C) Difference map of two single-condition maps. Dark: horizontal preferring. Light: vertical preferring. (D) Filtered map based on difference map, with both low and high pass (see text). (E) Following removal of large vessel pixels, the filtered map is clipped.

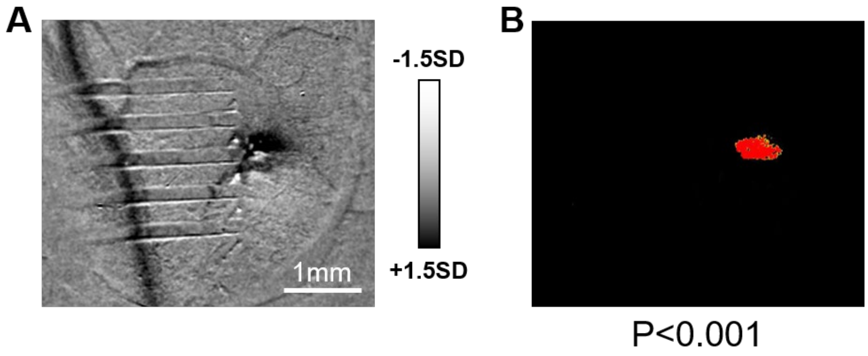


Supplementary Fig.3. The threshold used to choose activation region. (A) Original figure clipped with ±1.5SD (B) T-test map when $p<0.001$, which is the threshold we chose to indicate the activation region. The area is determined from the number of significant pixels and the diameter is the average of the long and short axes.

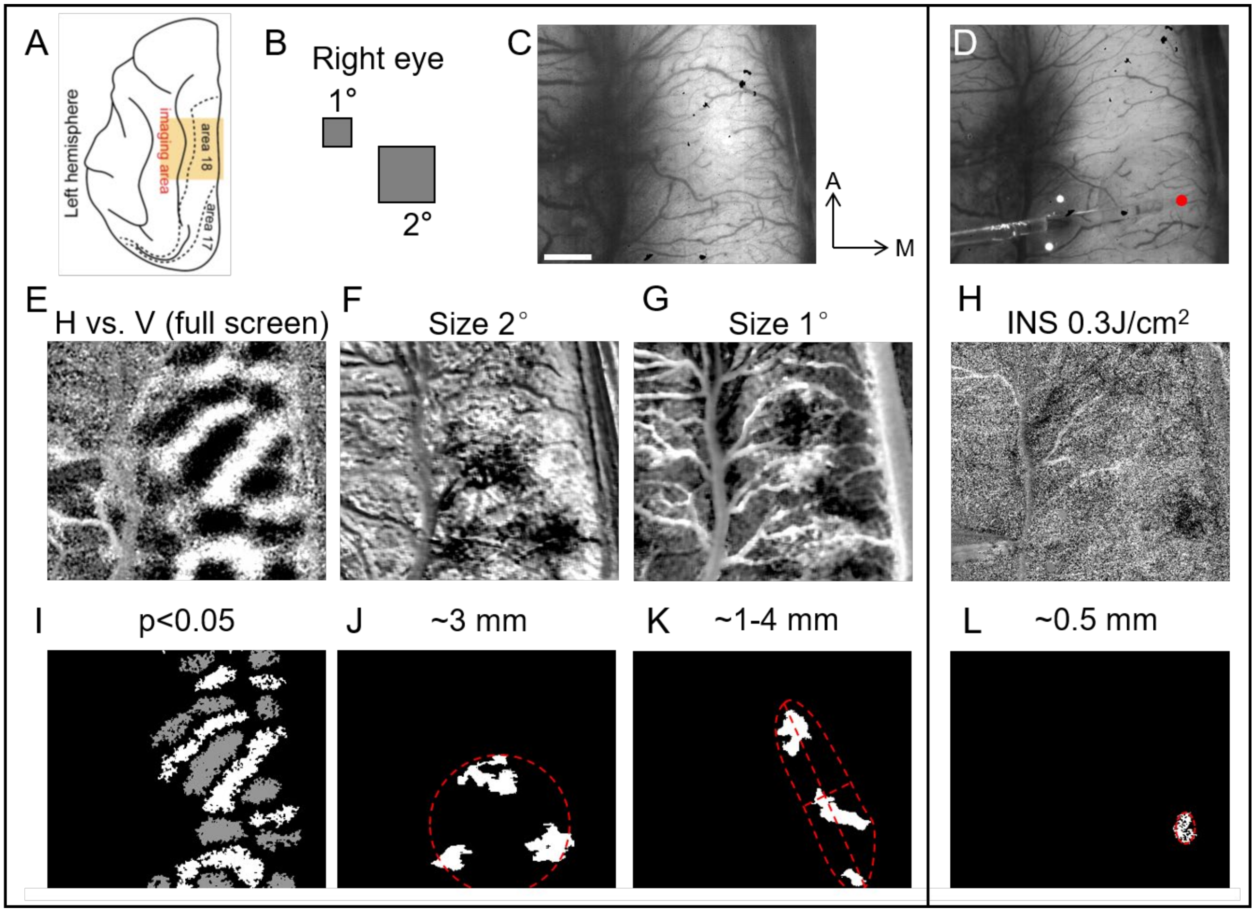


Supplementary Fig.4. Comparison of response size evoked by spot visual stimuli and single fiber INS stimulation. (A) Schematic picture of a cat cerebral cortex. Orange rectangle: approximate imaging field of view. Dotted lines: approximate areal borders. (B) Schematic of spot positions. 2° spot slightly more lateral than 1° spot. (C, D) Blood vessel maps recorded in the visual stimulation and INS conditions. (E, I) orientation map of 'H'-'V' full screen grating visual stimulation (binocular stimuli). (F, G) response map of 'vision'-'blank' from 2° (F), 1° (G) visual spot stimulation (monocular stimuli). (H) activation map in response to single fiber INS stimulation ('INS'-'blank', wavelength: 1870nm, frequency: 200Hz, radiant exposure: 0.3 $J \cdot cm^{-2}$, pulse width: 250us, pulse train: 0.5s). (J-L) significantly ($p<0.05$) activated pixels of F-H. Numbers at top of each panel: activation size of the cortex in the corresponding condition. Scale bar: 1mm. A: anterior, M: median.
