## Supplemental Methods for "A novel interface for cortical columnar neuromodulation with multi-point infrared neural stimulation"

### Temperature Rise induced by INS Stimulation

There have been multiple studies of temperature rise and damage thresholds of pulsed infrared neural stimulation (INS). Findings indicate that, for stimulation paradigms similar to that described here, thresholds, estimated from histological studies fall between 0.6J/cm$^2$ – 1J/cm$^2$ (rat cortex 0.4J/cm$^2$ and nonhuman primate cortex[1] ~0.6J/cm$^2$, human cortex[2] 0.6J/cm$^2$, human spinal roots[3] 1.09J/cm$^2$, cochlea 25uJ/pulse[4]).

Studies have also examined the temperature rise due to INS. Thompson et al.[5] modelled temperature rise in rat peripheral nerve and found that 250Hz stimulation with comparable parameters (200um fiber, pulse width 100usec, 1850nm, 25uJ) produced a temperature rise of 2.3°C. We have also addressed this issue using MRI thermometry (which measures the shift in proton resonance frequency caused by temperature increase). Using the same parameters used in this study for INS stimulation in ex vivo rat brains, MRI thermometry (spatial resolution: 1 mm isotropic voxels) reveals that the spatial extent of temperature increase is quite confined and the highest ΔT (measured at the strongest voxel in response to the highest intensity of 1J/cm2, red line) plateaus below 2°C (for optogenetic stimulation[6] see Luo et al. 2023).

To ensure the temperature rise of laser will not damage tissues, there is a In the thermometry study, we converted the direct proton resonance frequency phase shift data in acquired images to temperature change according to[6,7]:

$$\Delta T = \frac{\phi(T) - \phi(T_0)}{\gamma \alpha B_0 TE} \quad (1)$$

where $\phi(T)$ is the phase map at current time point, $\phi(T_0)$ is the phase map of the baseline image which is measured at room temperature (before INS), $\gamma$ here represents the gyromagnetic ratio of hydrogen (2.67 × $10^8$ rads per Tesla, constant), $\alpha$ is the PRF shift coefficient of water (-0.01 ppm per °C, constant), $B_0$ is the magnetic field strength (7 Tesla here), and TE is the echo time of imaging sequence (1.65 msec here). And in our condition, 1 °C ≅ 0.031 rads.

The temperature of the voxel with greatest phase shift at the location of the optic fiber tip was selected, and the temperature increase in the 9 isolated measurements was averaged for further analysis. ANOVA and paired-sample t-test were conducted to statistically analyze the temperature changes before and after INS, as well as the differences across different power intensities at the same time point. The data was analyzed using MATLAB.

In sum, assessments of heat induced damage via histological methods, thermometry, and modelling, all indicate that INS delivered with these parameters are non-damaging. Note also that, based on these studies, we have conducted INS in human cortex[2] and there are currently clinical trials using INS (cochlea: Richter NCT05110183, peripheral nerves: Jansen NCT04601337).
